# Meiofaunal communities flourish in Antarctic marine sediments despite the harsh environmental conditions

**DOI:** 10.64898/2026.05.19.726228

**Authors:** Marta García-Cobo, Diego Fontaneto, Ester M. Eckert, Raffaella Sabatino, Matteo Cecchetto, Stefano Schiaparelli, Alejandro Martínez

**Affiliations:** Universidad Complutense de Madrid, Faculty of Biological Sciences, Department of Biodiversity, Ecology and Evolution. C/ José Antonio Novais 12. 28040 Madrid, Spain; Molecular Ecology Group (MEG), Water Research Institute (IRSA), National Research Council of Italy (CNR). Largo Tonolli, 50. 28922, Verbania, Italy; Department of Earth, Environmental and Life Sciences (DISTAV), University of Genoa, Corso Europa 26, Genoa, Italy, Department of Earth, Environmental and Life Sciences (DISTAV), University of Genoa, Corso Europa 26, Genoa, Italy; Italian National Antarctic Museum (MNA, Section of Genoa), University of Genoa, Viale Benedetto XV No. 5, Genoa, Italy, Italian National Antarctic Museum (MNA, Section of Genoa), University of Genoa, Viale Benedetto XV 5, Genoa, Italy

## Abstract

While Antarctic terrestrial ecosystems support low metazoan diversity, the surrounding marine macrobenthos is rich. However, marine meiofauna remains historically neglected, leaving its diversity patterns unclear. In this study, we used 18S rRNA gene metabarcoding alongside an enhanced taxonomic annotation pipeline to characterize marine meiofauna diversity in the Ross Sea, comparing it to global datasets. We evaluated how depth, habitat type, and mesh size influence community structures to test if habitat heterogeneity drives diversity despite the harsh Southern Ocean conditions. Our results revealed exceptionally high diversity, with metazoans richness comparable to or higher than temperate regions. Although environmental variables had limited effects on taxonomic richness, they significantly shaped community composition, with habitat type explaining the highest proportion of variance. Interestingly, we detected several ASVs 100% identical to North Sea and North Atlantic sequences, likely reflecting the limited taxonomic resolution of the 18S marker rather than global dispersal (the “meiofaunal paradox”). Overall, these findings demonstrate that Antarctic marine sediments host rich meiofaunal communities where ecological processes operate similarly to other global regions, contrasting sharply with depauperate continental Antarctic ecosystems.

## INTRODUCTION

Antarctica, often referred to as the “white continent”, is known to have among the most challenging conditions for life on Earth (Gonzalez & Vasallo, 2019). Approximately 98% of its surface is permanently covered by ice and snow, with persistent subzero temperatures, intense desiccation, strong winds and pronounced seasonal fluctuations in light availability, which directly influence primary production (Convey et al., 2010, Clarke & Harris, 2003). In addition, Antarctica’s nutrient-poor substrates and long-standing geographic isolation add further constraints on biological colonization and survival. As a result, Antarctic terrestrial environments support very low metazoan diversity compared to other continents, leading to depauperate terrestrial biota (Convey et al. 2008; Pertierra et al., 2025). To date, approximately 400 terrestrial and freshwater metazoan species have been recorded in continental Antarctica (Convey, 2010; Dartnall, 2017; Pertierra et al., 2024, 2025), a very small number compared to other biogeographical regions. This pattern is not due to poor knowledge, but reflects a biological reality (Pertierra et al., 2025). Limno-terrestrial meiofauna—defined in ecological studies as metazoans retained between 1 - 0.030 mm mesh sizes (Martínez et al. 2025)—are among the most successful colonizers of ice-free areas. Yet, Antarctic limno-terrestrial meiofauna, which represents most of the fauna, is composed only by few species of nematodes, tardigrades, rotifers, collembolans, mites, and flatworms, typically forming simple communities with a handful of species (Freckman & Virginia, 1997; Andrássy & Gibson, 2006; Pertierra et al., 2024, 2025).

In contrast to the harsh and species-poor terrestrial Antarctic habitats (Convey et al., 2014a; Dartnall et al., 2017), the surrounding marine ecosystems seem to exhibit relatively higher biodiversity. The Southern Ocean, which encircles Antarctica, acts as a thermal and chemical buffer, moderating environmental extremes through its high heat capacity, solvent properties, and stable composition (Clarke & Harris, 2003). To date, more than 17,000 metazoan species have been documented in the Southern Ocean (De Broyer et al., 2014; Peck, 2018), though this diversity is likely underestimated due to the unbalance sampling effort across different taxa and habitats (Brandt et al., 2007; Kaiser et al., 2013). Nevertheless, Antarctic metazoan marine species richness is generally comparable to the global averages, and even exceeds them for certain groups such as pycnogonids, which diversified in the Southern Ocean (Barnes & Peck, 2008). Still, also Antarctic marine biodiversity can reflect a reduced representation of global diversity, as happens for fish, with very few species, represented mostly by a radiation of one group, the Notothenioidei and species from two other families, Liparidae and Xoarcidae (Matschiner et al., 2011; Di Prisco et al., 2012). Although representing 10% of the world’s ocean, the Southern Ocean hosts only 1% of the global diversity of fish (Di Prisco et al., 2012). In general, historical legacy has played a great role in shaping present Antarctic assemblages that largely correspond to those that survived glacial cycles (Convey et al. 2014b).

Marine meiofauna represents a substantial but historically neglected component of Antarctic marine biodiversity (Ingels et al. 2023). Early studies based on morphological identifications revealed a higher meiofaunal diversity in marine sediments in comparison to poor terrestrial environments, encompassing nematodes (Cobb, 1914; 1930; Lee et al., 2017), copepods (Apostolov & Pandourski, 1999), annelids (Rota & Carchini, 1999), platyhelminthes (Artois et al. 2000; Volonterio & Ponce de León, 2018), kinorhynchs (Sørensen, 2008; Sørensen et al., 2025a), gastrotrichs (Kieneke, 2010; Sørensen et al., 2025), and tardigrades (Villora-Moreno, 1999). Despite this taxonomic diversity, community-level studies remain scarce and have largely focused on nematodes (Vanhove et al., 1998; Hong et al., 2011) or on responses to specific environmental disturbances such as iceberg scouring, pollution, or the establishment of marine protected areas (Lee et al., 2001; Stark et al., 2017; Semprucci et al., 2021; França et al., 2024; De Araújo França et al., 2026). This research gap likely stems not only from the inherent challenges of meiofaunal work (Fonseca et al., 2018; Martínez et al., 2025), but also from the logistical constraints of conducting fieldwork in remote Antarctic marine environments (Clarke & Harris, 2003). As a result, our understanding of Antarctic meiofaunal diversity lags behind that of other biogeographical regions; an unfortunate shortfall, given Antarctica’s importance for assessing ecosystem resilience in the face of climate change (Jeunen et al., 2023).

Molecular tools such as DNA metabarcoding are increasingly bridging these knowledge gaps. Pioneering studies suggested that the diversity of Antarctic marine sediments may not be depauperated, both in terms of taxonomic and phylogenetic diversity richness (Fonseca et al., 2017; Clarke et al., 2021). These studies also revealed that previous morphology-based surveys had captured only a fraction of the actual diversity, which includes a combination of both widely distributed and locally endemic marine Antarctic species (Brannock et al., 2018). More recently, multigene approaches have been used to explore broader patterns of diversity (Fonseca et al., 2022). However, these studies typically used total marine metazoan richness as a response variable, without distinguishing between permanent meiofauna and transient forms such as larvae, juveniles, or body fragments (collectively referred to as temporary or accidental meiofauna) (Martínez et al. 2025).

The goal of this study is to test whether marine meiofauna in Antarctica, contrary to the very poor freshwater and limno-terrestrial Antarctic meiofauna, is rich and not depleted. To achieve this, we focused on metabarcoding of several samples, developed an enhanced taxonomic annotation pipeline, in addition to assigning each molecular taxonomic unit (Amplicon Sequence Variant, ASV, in our case) to functional groups: permanent meiofauna, temporary/accidental meiofauna, and parasites. Furthermore, samples were fractioned using a stack of sieves with different mesh sizes to assess how mesh selection influences taxonomic resolution. Specifically, we examined overall diversity across taxonomic groups of permanent marine meiofauna, in addition to assessing if ecological differences between habitat and depth gradients could explain the locally observed patterns. Finally, we qualitatively compared the results of our survey with similar surveys in other parts of the world and assessed the proportions of ASVs lacking clear taxonomic assignment at the genus or family level to reflect the degree of unexplored or undescribed diversity in Antarctic benthic meiofaunal communities.

## MATERIALS AND METHODS

### Data collection

We collected 42 samples from 17 sites from the Ross Sea area adjacent to Victoria Land in Antarctica (Figure 1), covering six types of habitats: epilithic, organic, spicule, gravel, sand, and silt. Sampling site locations and characteristics are reported in Table 1. Each sample consisted of about half square meter of sediment collected with a grab operated from a boat. Each sample was filtered through a sieve of 1 mm mesh size to remove larger particles and macrofauna, and then, the filtrate was washed through four nested sieves of 200, 100, 50 and 20 µm mesh size. The organisms that were retained in each sieve were collected and stored in 50 ml tubes in ethanol 96% in a freezer at -20 °C.

**Table 1.** List of sampling sites, including the name of the sites, coordinates, the number of samples, the name of each sample, the number of subsamples obtained after washing the sediment through different mesh sizes (20, 50, 100 and 200 µm), sampling depth and type of habitat.

| site (map) | longitude | latitude | number<br>of<br>samples | name of samples<br>(number of<br>subsamples) | depth (m) | habitat |
| --- | --- | --- | --- | --- | --- | --- |
| ADA I | 164,1268 | -74,6983 | 2 | ADA 1-1 (4) | 40 | sand |
|  |  |  |  | ADA 1-3 (4) | 40 | sand |
| ADA II | 164,1222 | -74,6968 | 2 | ADA 1-4 (4) | 70 | silt |
|  |  |  |  | ADA 1-5 (4) | 19 | gravel |
| ADCO I | 163,9897 | -74,7719 | 2 | ADCO 1 (4) | 33 | epilithic |
|  |  |  |  | ADCO 2 (4) | 33 | silt |
| ADCO II | 164,0004 | -74,7721 | 1 | ADCO 3 (4) | 38 | silt |
| ADCO III | 164,0014 | -74,7703 | 1 | ADCO 4 (4) | 58 | epilithic |
| ADCO IV | 164,03 | -74,7695 | 2 | ADCO 6 (4) | 70 | silt |
|  |  |  |  | ADCO 8 (4) | 20 | sand |
| **AMORPHOUS** | 164,0366 | -74,6872 | 6 | AMOURPHOUS 1 A (4) | 20 | sand |
|  |  |  |  | AMOURPHOUS 1 B (4) | 20 | sand |
|  |  |  |  | AMOURPHOUS 2 (4) | 20 | sand |
|  |  |  |  | AMOURPHOUS 3 (4) | 20 | sand |
|  |  |  |  | AMOURPHOUS 4 (4) | 20 | gravel |
|  |  |  |  | AMOURPHOUS 5 (4) | 20 | sand |
| **CAL** | 164,0911 | -74,7519 | 4 | CAL 1 (4) | 50 | silt |
|  |  |  |  | CAL 2A (4) | 65 | organic |
|  |  |  |  | CAL 2B (4) | 65 | organic |
|  |  |  |  | CAL 3 (4) | 20 | silt |
| **FAR I** | 164,121 | -74,7186 | 1 | FAR 1 (4) | 40 | sand |
| **FAR II** | 164,1128 | -74,7186 | 1 | FAR 2 (4) | 19 | sand |
| **MAREOGR I** | 164,1168 | -74,6917 | 1 | MAREOGR 1 A (4) | 25 | sand |
|  |  |  |  | MAREOGR 1 B (4) | 25 | sand |
| **MAREOGR II** | 164,1261 | -74,7352 | 1 | MAREOGR 2 (4) | 49 | sand |
| **MOL** | 164,0469 | -74,7342 | 1 | MOL 1 (4) | 20 | gravel |
| **PN** | 164,0695 | -74,6756 | 7 | PN 1A (4) | 24 | epilithic |
|  |  |  |  | PN 1B (4) | 24 | epilithic |
|  |  |  |  | PN 1C (4) | 24 | epilithic |
|  |  |  |  | PN 2 (4) | 24 | gravel |
|  |  |  |  | PN 3 (4) | 24 | sand |
|  |  |  |  | PN 4 (4) | 24 | gravel |
|  |  |  |  | PN 5 (4) | 24 | gravel |
| **SPIAGG** | 164,042 | -74,7011 | 5 | SPIAGG 1 (4) | 22 | silt |
|  |  |  |  | SPIAGG 2 (4) | 22 | silt |
|  |  |  |  | SPIAGG 3 (4) | 22 | silt |
|  |  |  |  | SPIAGG 4 (4) | 22 | silt |
|  |  |  |  | SPIAGG 5 (4) | 22 | silt |
| **SPIAGGIA MOLO** | 164,042 | -74,7011 | 1 | SPIAGGIA MOLO 1 (4) | 0 | sand |
| **TETH** | 164,042 | -74,6989 | 4 | TETH 1 (4) | 55 | silt |
|  |  |  |  | TETH 2 (3) | 70 | silt |
|  |  |  |  | TETH 5 (4) | 70 | spicule |
|  |  |  |  | TETH 6 (2) | 70 | spicule |
| Total: | 17 |  | 42 | 169 subsamples |  |  |
| sites |  |  | samples |  |  |  |

**Figure 1.**
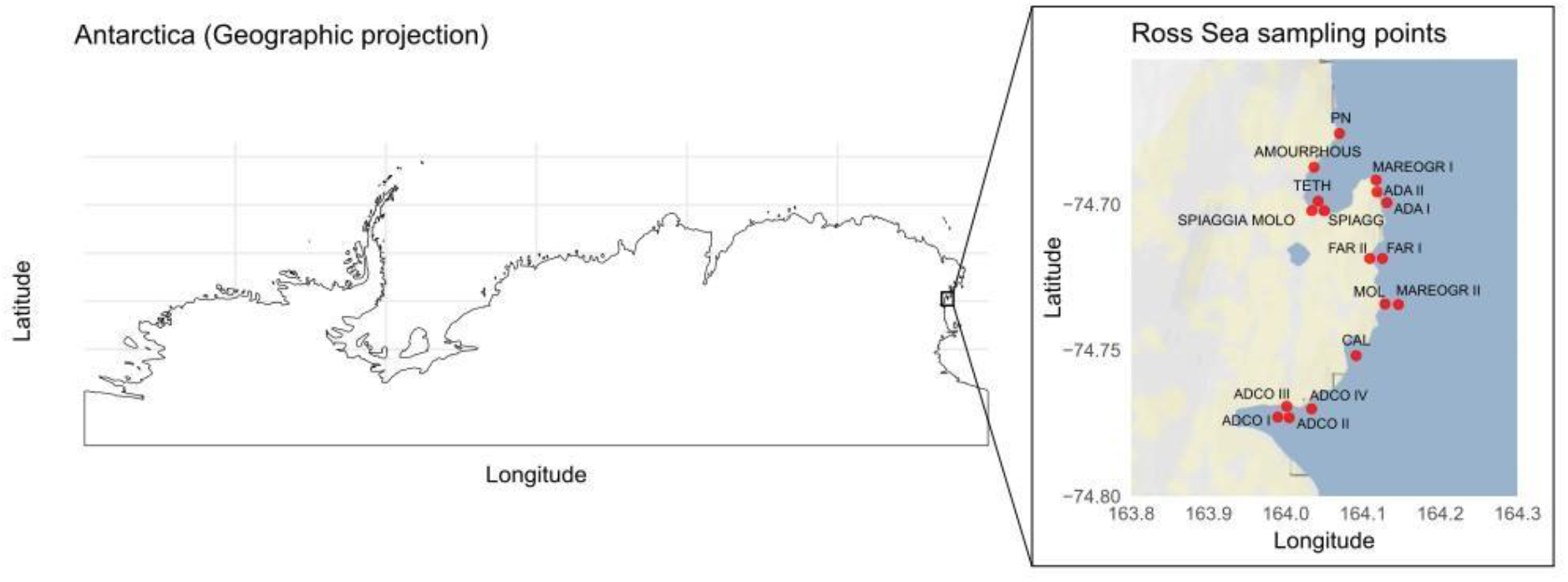
Map of Antarctica showing the 17 sampled sites in the Ross Sea.

Sampling locations were plotted with the R version 4.5.1 (R Core Team, 2025) on high-resolution maps of the Ross Sea obtained from the “ggmap” package version 4.0.0 (Kahle & Wickham, 2019) and customized with “ggplot2” version 3.5.1 (Wickham, 2016). An overview map of Antarctica was created using the “sf” version 1.6.5 and “rnaturalearth” version 1.0.1 packages (Pebesma & Bivand, 2023; Massicotte & South, 2023).

### DNA extraction and sequencing

Samples were processed for DNA extraction and sequencing as previously described (Sterrer et al., 2022). Briefly, an aliquot of the ethanol-preserved samples was dried under a laminar flow hood and treated with a Proteinase K solution for 2 h at 56 °C. DNA was then extracted using the commercial DNeasy PowerSoil extraction kit (Qiagen) and sent to an external company (IGA Technology, Udine, Italy) for sequencing of the V1–V2 hypervariable regions of the 18S rRNA gene. Raw sequences were made publicly available in NCBI under the project accession number XXX.

### Bioinformatic analyses

To accurately characterize diversity patterns associated with Antarctic meiofauna, we implemented an enhanced annotation pipeline tailored to correctly classify meiofauna in different taxa and functional groups. Raw sequences were first imported into R and processed using the DADA2 pipeline, following the online tutorial (Callahan et al., 2016). Briefly, sequences were quality-filtered and trimmed. Then, sequences were merged in a 6-step annotation process. (1) ASVs were initially classified using the ‘classify.seqs’ function and the SILVA v138.1 database to obtain a coarse taxonomic annotation. (2) Annotated sequences were aligned using the ‘e-INS-i’ algorithm in “MAFFT” software version 7 (Katoh & Toh, 2010). A pairwise distance matrix and neighbor-joining tree were generated using ‘dist.dna’ (with pairwise.deletion = TRUE) and ‘njs’ functions from the “ape” R package v.5.8.1 (Paradis & Schliep, 2009). Clades containing at least one metazoan ASV were identified, and all ASVs contained in the putative metazoan clades were extracted using ‘getMRCA’ and ‘extract.clade’ functions from the “ape” R package. (3) Those “metazoan candidate ASVs” were then BLASTed against a curated 18S rRNA reference database. BLAST was run locally, and the top 10 hits for each ASV were retained and summarized for taxonomic inference. (4) All ASVs assigned to major metazoan groups by BLAST were then aligned alongside reference sequences and subjected to group-specific neighbor-joining tree reconstruction. ASVs that clustered within the ingroup were retained, whereas those clustering with outgroup sequences were flagged as doubtful and excluded from further analyses. (5) A summary table was compiled including annotations from SILVA, BLAST, and neighbor-joining analyses. Each ASV was assigned to a taxonomic group: the group corresponded to the phylum (e.g. Annelida, Bryozoa, Mollusca, etc.), except for Arthropoda, which were separated into multiple groups, including Acari, Branchiopoda, Copepoda, Cumacea, Decapoda, Euphasiacea, Hexapoda, Isopoda, and Ostracoda. (6) Assessment of taxonomic novelty. For each unique ASV, we recorded the percentage of sequence identity from the top BLAST hit against the reference database. To assess the degree of novelty, ASVs were categorized into similarity tiers based on established 18SrRNA gene thresholds: values <95% were used to identify deep-branching lineages potentially representing potentially undescribed families or higher ranks (Caron et al., 2009; Pawlowski et al., 2012).

Each ASV was also assigned to one of three ecological categories based on their best matches against the reference library: (i) permanent meiofauna, (ii) temporary/accidental meiofauna, or (iii) parasites. Here, temporary meiofauna refers not only to organisms with a transient meiofaunal stage in their life cycle but also to fragmented remains of macrofaunal species—both indistinguishable in a metabarcoding context. In addition, Rotifera and Acanthocephala were kept separated and not included as Syndermata (Vasilikopoulos et al., 2024), to facilitate the separation of the strictly parasitic acanthocephalans.

To enhance comparability of results with diversity from other biogeographical areas, the same procedure was performed on all the raw data of the previously published datasets dealing with metabarcoding marine meiofauna using the same marker that we could find in the literature.

### Statistical analyses

All R scripts and datasets used for data wrangling, statistical analyses, and visualization are hosted in a public GitHub repository /r/Antarctica-metabarcoding-meiofauna-4C26/.

First, we qualitatively compared the results of our survey with other metabarcoding studies on marine meiofauna in other biogeographical regions. Then, to assess the detailed drivers of meiofauna diversity in Antarctic sediments, we quantitatively analyzed the effects of habitat type and sampling depth on both alpha and beta diversity, while accounting for mesh size as a potential influencing factor. ASVs identifying temporary meiofauna or parasites were discarded from all analyses to account only for the actual permanent meiofaunal diversity.

#### Comparison with metabarcoding studies

To contextualize our findings within the broader framework of global marine biodiversity, we performed a comparative analysis between our dataset and all previously published 18S rRNA gene metabarcoding surveys of marine meiofauna that used similar metabarcoding approaches. We first performed a standardized Google Scholar search using the keywords “meiofauna”, “NGS”, and “metabarcoding”. We then screened the titles and abstracts of all retrieved papers and retained studies targeting the V1–V2 region of the 18S rRNA gene using Illumina sequencing. Of the 22 retrieved papers, 11 had sequence data publicly available, whereas for 6 studies we obtained the raw sequences directly from the authors upon reasonable request. From this dataset, three studies were excluded because of poor-quality reverse reads, two because they included data from the deep sea (Brandt et al., 2021; Klunder et al., 2020) and one because it dealt with mock communities (Rossel et al., 2019), leaving us with 10 studies. Exceptionally, these 10 studies also include Fonseca et al. (2017), despite its use of pyrosequencing, because it represents the only comparable study conducted in Antarctica. The annotated summary of all downloaded papers, along with the Bioproject accession numbers, and comments on the selection process are summarized in Table S7.

We then analysed raw DNA sequence data through the same pipeline that we used for our survey, in order to identify ASVs with the same approach, allowing comparability. This allowed us to calculate taxonomic richness, community composition and taxonomic novelty using the same criteria across all datasets, regardless of the different analytical approaches and pipelines in different studies.

Connectivity patterns were visualized through geospatial network analysis using the ‘tbl_graph’ and ‘ggraph’ functions from the packages “tidygraph” version 1.3.1 and “ggraph” version 2.2.2 (Pedersen, 2024 and Pedersen, 2025, respectively). Nodes were defined by the geographic coordinates of each study, while edges represented shared ASVs. We implemented a multi-threat approach using the ‘uncount’ function from the package “tidyr” version 1.3.1 (Wickham et al., 2024) to replicate edges based on their weight, followed by visualization with the ‘geom_edge_fan’ function from the aforementioned package “ggraph”. Furthermore, taxonomic composition across studies was compared using a normalized heatmap, in which the number of ASVs of each taxonomic group was divided by the total number of ASVs reads in each study. The resulting matrix was log-transformed and visualized using the ‘geom_tile’ and ‘geom_text’ function from the “ggplot2” package version 4.0.0 (Wickham, 2016), allowing for a standardized comparison of the relative importance of each meiofaunal group across different regions. For the network analysis, references representing the same sampling surveys or geographical location (Haenel et al., 2017; Atherton & Jondelius, 2020) were consolidated into a single node to prevent artificial inflation of connectivity due to redundant sampling. Geographic coordinates for consolidated nodes were calculated as the mean position of the original sampling sites.

#### Alpha diversity

Alpha diversity was measured for taxonomic diversity as the number of ASVs. Phylogenetic diversity was measured as the total branch length of the tree for the species present in each unit of diversity, using the function ‘alpha’ from the “BAT” package version 2.11.0 (Cardoso et al., 2025). The phylogenetic tree was calculated from the alignment of unique ASVs with the function ‘AlignSeqs’ from the package “DECIPHER” version 3.4.0 (Wright, 2016), computing a maximum-likelihood distance matrix according to the function ‘dist.ml’ of the package “phangorn” version 2.12.1 (Schliep, 2011) and the reconstruction of a Neighbor-Joining (NJ) tree.

To evaluate differences in taxonomic and phylogenetic diversity, we fitted Generalized Linear Mixed Models (GLMM) using the function ‘glmmTMB’ in the “glmmTMB” package version 1.1.10 (Brooks et al., 2017). For taxonomic diversity, Poisson distributions were used unless overdispersion was detected using the ‘check_overdispersion’ function from the “performance” package version 0.13.0 (Lüdecke et al., 2021). In cases of overdispersion, negative binomial distributions were used to account for overdispersed count data. For phylogenetic diversity, we used a Tweedie distribution with a log-link function, which is particularly well-suited for continuous positive data that may include zeros or exhibit high variance (Shono, 2008). In both types of models, fixed effects included sampling depth, habitat type and mesh size, with sample ID as a random effect to account for the hierarchical structure of the data and (log transformed) read number as an offset to account for the potential bias due to differences in sequencing depth. Model performance and assumptions (e.g. normality of residuals, homoscedasticity, collinearity, outliers, etc.) were verified using the ‘check_model’ function of the “performance” package version 0.13.0 (Lüdecke et al., 2021). Significance of fixed categorical effects was tested via Type II Wald chi-square tests using the ‘Anova’ function of the “car” package version 3.1.3 (Fox & Weisberg, 2019) and the summary of the model. For categorical variables (habitat and mesh size), post-hoc pairwise comparisons of Estimated Marginal Means (EMMs) were performed with Tukey’s adjustment using the function ‘emmeans’ of the package “emmeans” version 1.10.7 (Lenth, 2025). To visualize the partial effects of mesh size and habitat type on ASV richness, we generated predicted value plots using Estimated Marginal Means (EMMs) derived from the GLMMs. Predictions and their associated 95% Wald confidence intervals were computed by holding continuous covariates at their mean and averaging across categorical factors.

#### Beta diversity

Differences in community taxonomic composition were assessed using the Jaccard dissimilarity index for presence/absence community matrixes. Phylogenetic beta diversity was calculated by incorporating the previously reconstructed Neighbor-Joining tree using the ‘beta’ function from the “BAT” package version 2.11.0 (Cardoso et al., 2025). The two types of beta diversity, taxonomic and phylogenetic, were also partitioned into their turnover and nestedness components.

To test the influence of habitat, depth, and mesh size on beta diversity, we performed Permutational Multivariate Analysis of Variance (PERMANOVA) using the function ‘adonis2’ from the “vegan” package version 2.6.10. The models included the log transformed total number of reads as a covariate to control for sequencing effort. Permutations were restricted within each sample ID using the *strata* argument to account for the dependency between subsamples (different meshes) from the same sample. Furthermore, significance was assessed using the *by=“margin”* parameter, ensuring that the marginal effect of each term was evaluated independently of its order in the model.

All aforementioned analyses were repeated for the total meiofaunal community and separately for those taxonomic groups that were sufficiently represented across samples. Rare or low-frequency taxonomic groups were excluded from individual statistical modelling to avoid low statistical power and model convergence issues, although they were included in all community-level (total meiofauna) analyses.

Venn diagram was drawn with the ‘venn.diagram’ function from the “VennDiagram” package version 1.7.3 (Chen, 2022) to illustrate the number of exclusive and overlapping ASVs among the different mesh sizes.

#### Taxonomic novelty

From the proportion of ASVs with a BLAST hit below 95%, representing potentially undescribed families or higher ranks (Caron et al., 2009; Pawlowski et al., 2012), we fitted a normal distribution. We then estimated mean, standard deviation, and assessed whether the observed proportion from Antarctica was significantly different from the proportions of the previously published surveys in other biogeographical regions.

## RESULTS

We obtained 2329 metazoan Amplicon Sequence Variants (ASVs) in this study, corresponding to the following 21 phyla: Arthropoda (including the groups Copepoda (1114 ASVs), Ostracoda (173), Acari (9), Branchiopoda (6), Isopoda (5), Decapoda (4), Cumacea (1), Euphasiacea (1), and Hexapoda (1)), Nematoda (937), Annelida (298), Platyhelminthes (193), Cnidaria (161), Echinodermata (85), Gastrotricha (82), Acanthocephala (65), Porifera (54), Mollusca (48), Gnathostomulida (41), Xenacoelomorpha (34), Nemertea (31), Rotifera (21), Tardigrada (16), Bryozoa (9), Tunicata (9), Chordata (5), Brachiopoda (4), Entoprocta (1), and Priapulida (1). After filtering out 74 parasites ASVs and 681 temporary meiofauna ASVs, a total of 1793 ASVs were identified as permanent meiofauna.

The permanent meiofauna of our study comprised 10 phyla: Arthropoda (with the groups: Copepoda (1114 ASVs), Ostracoda (173), Acari (9), Branchiopoda (5), and Isopoda (1)), Nematoda (937), Platyhelminthes (177), Gastrotricha (82), Gnathostomulida (41), Annelida (40), Xenacoelomorpha (34), Rotifera (21), Tardigrada (16), and Nemertea (7) (Figure 2). In comparison to a previous metabarcoding study carried out in Antarctica (Fonseca et al., 2017), our survey added gnathostomulids (already mentioned in Sterrer et al., 2022), nemerteans, tardigrades, and xenacoelomorphs but did not find any kinorhynchs. Using the same taxonomic groups (equal to phyla except for further divisions for arthropods), Antarctic diversity was comparable to, if not even higher than, that of other studies carried out at different latitudes using the same fragment of 18S rRNA gene, both for the number of major taxonomic groups and for the number of ASVs, even if the large differences in sequencing depth between studies make the comparisons qualitative (Figure 2). The comparison included disparate areas like the coast of Massachusetts (Polinski, 2019), the Asinara National Park, Italy (Martínet at al., 2020), the North Sea (Cordier et al., 2021; Degenhart et al., 2021; Haenel et al., 2017; Atherton & Jondelious, 2020), the Baltic Sea (Kapshyna et al., 2024), and the Arctic (Mazukiewicz et al., 2024) (Figure 2; Table S1).

**Figure 2.**
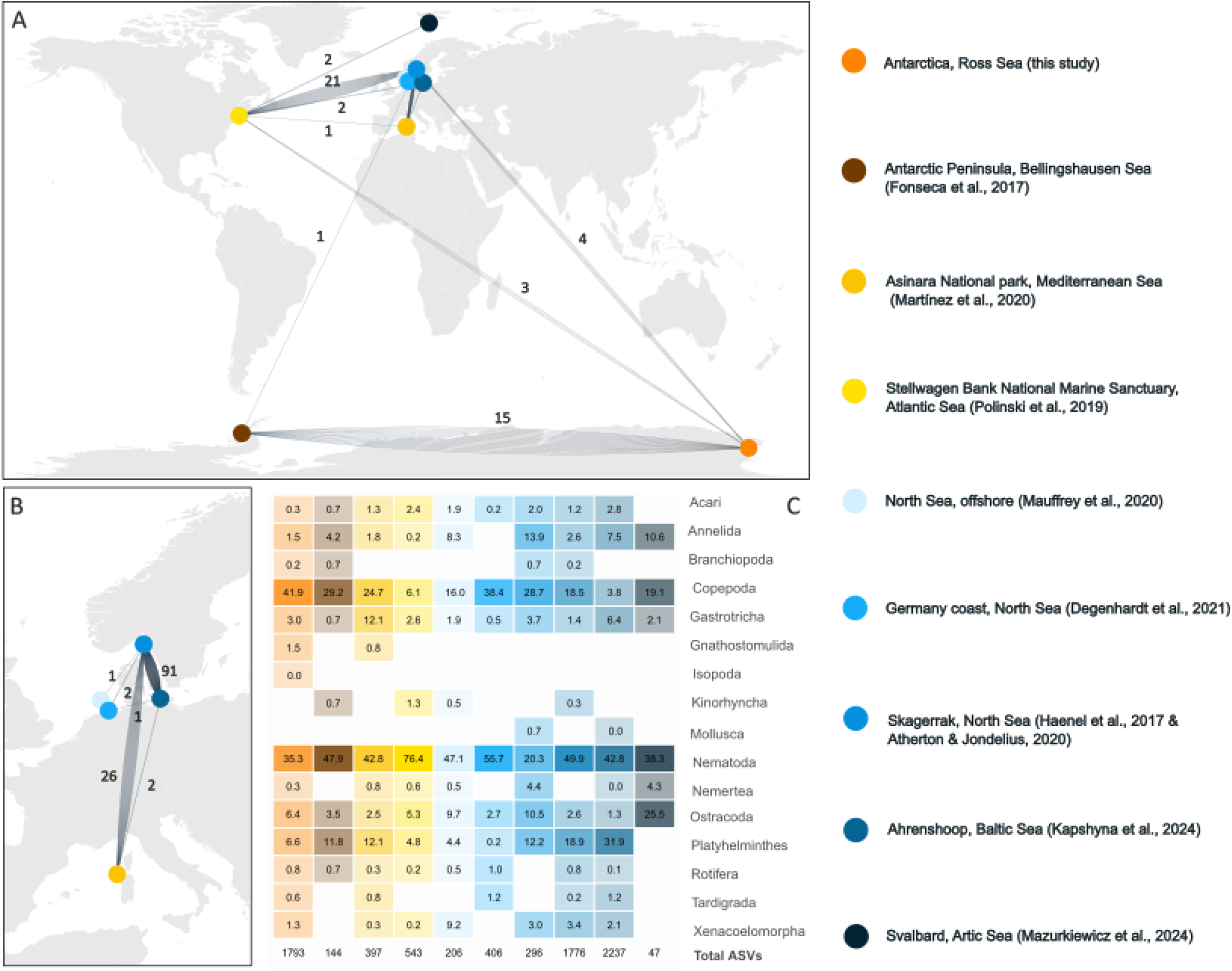
Comparative analysis of ASV sharing and taxonomic distribution across studies. (A, B) Network analyses illustrate the number of shared ASVs among the included metabarcoding datasets, (A) worldwide and (B) focusing on Europe, as it contains multiple studies; nodes represent individual studies and edge thickness corresponds to the number of shared sequences, written close to each edge. (C) Heatmap showing the proportion (%) of ASVs for each phylum, with arthropods divided in further taxonomic groups. The color scale for the heatmap corresponds to the study-specific identifiers (colored dots) shown in the legend to the right. Note that proportions <0.1% appear as 0.0 as values are rounded to a single decimal.

Regarding shared exact sequences, our study identified 15 ASVs in common with the only other available Antarctic survey (Fonseca et al., 2017). These included copepods (Canthocamptidae and Ectinosomatidae), gastrotrichs (Macrodasydae), nematodes (Leptolaimidae, Chromadoridae, Anticomidae and Xyalidae), platyhelminthes (Dolichomacrostomidae, Koinocystidae, and Polycystidae) and annelids (Enchytraeidae and Naididae). Furthermore, our Antarctic samples shared three nematode (Monhysteridae) ASVs with a metabarcoding study from the Atlantic Ocean off Massachusetts (Polinski et al., 2019), a single copepode ASV (Harpacticoida, Ectinosomatidae) from the North Sea (Haenel, 2017), and three ASVs with another North Sea survey by Atherton and Jondelius (2020): one Oithonidae (Copepoda), as well as a Synchaetidae and a Dicranophoridae (Rotifera).

### Alpha diversity: taxonomic and phylogenetic richness

For the total meiofauna community, none of the variables significantly affected overall phylogenetic, whereas taxonomic richness was only significantly affected by the habitat type (GLMM: *χ*^2^=12.293, p=0.031) (Table 2 & Table 3).

**Table 2.** Results of the Analysis of Deviance (Type II Wald *χ*^2^ tests) for the Generalized Linear Mixed Models evaluating the effects of the variables–sampling depth, mesh size and type of habitat–on alpha taxonomic diversity of the total meiofauna and by each meiofaunal taxa (Note: only taxa with sufficient abundance and occurrence across samples were modelled to ensure statistical power and model convergence).

|  |  | Meiofauna | Copepoda | Nematoda | Platyhelminthes | Ostracoda | Annelida | Gastrot richa | Xenacoelomorpha |
| --- | --- | --- | --- | --- | --- | --- | --- | --- | --- |
| depth (m) | Est. | 0,005 | -0,037 | 0,109 | -0,463 | -0,068 | -0,519 | 0,159 | 0,277 |
| | $\chi^2$ | 0,015 | 0,344 | 3,17 | 5,751 | 0,164 | 0,906 | 0,472 | 0,694 |
|  | df | 1 | 1 | 1 | 1 | 1 | 1 | 1 | 1 |
|  | p | 0,903 | 0,558 | 0,075 | **0,016** | 0,685 | 0,341 | 0,492 | 0,405 |
| mesh size | $\chi^2$ | 3,283 | 4,286 | 4,877 | 3,865 | 1,147 | 1,15 | 11,456 | 5,899 |
|  | df | 3 | 3 | 3 | 3 | 3 | 3 | 3 | 3 |
|  | p | 0,350 | 0,232 | 0,181 | 0,276 | 0,766 | 0,765 | **0,010** | 0,117 |
| habitat | $\chi^2$ | 12,293 | 12,077 | 12,847 | 15,141 | 13,81 | 8,708 | 2,979 | 1,704 |
|  | df | 5 | 5 | 5 | 5 | 5 | 5 | 5 | 5 |
|  | p | **0,031** | **0,034** | **0,025** | **0,010** | **0,017** | 0,121 | 0,703 | 0,888 |
| Random effects | N | 161 | 161 | 161 | 161 | 161 | 161 | 161 | 161 |
|  | R<sup>2</sup> Marginal/Conditional | 0,002 / 0,003 | - | 0,005 / 0,007 | - | 0,064 / 0,071 | - | - | 0,834 / 0,852 |

**Table 3.** Results of the Analysis of Deviance (Type II Wald *χ*^2^ tests on) for the Generalized Linear Mixed Models evaluating the effects of the variables–sampling depth, mesh size and type of habitat– on alpha phylogenetic diversity of the total meiofauna and by each meiofaunal taxa (Note: only taxa with sufficient abundance and occurrence across samples were modelled to ensure statistical power and model convergence).

|  |  | Meiofauna | Copepod | Nematod | Platyhelminthes | Ostracod | Gastrot rich |
| --- | --- | --- | --- | --- | --- | --- | --- |
|  |  | a | a | a | es | a | a |
| depth (m) | Est. | -0,048 | -0,069 | 0,038 | -0,637 | -0,476 | 0,085 |
| | $\chi^2$ | 0,439 | 0,592 | 0,201 | 8,541 | 5,419 | 0,094 |
|  | df | 1 | 1 | 1 | 1 | 1 | 1 |
|  | p | 0,508 | 0,441 | 0,654 | **0,003** | **0,02** | 0,759 |
| mesh size | $\chi^2$ | 4,873 | 1,14 | 4,231 | 0,379 | 2,943 | 9,171 |
|  | df | 3 | 3 | 3 | 3 | 3 | 3 |
|  | p | 0,181 | 0,767 | 0,238 | 0,945 | 0,401 | **0,027** |
| habitat | $\chi^2$ | 7,04 | 4,016 | 7,184 | 11,694 | 16,678 | 2,852 |
|  | df | 5 | 5 | 5 | 5 | 5 | 5 |
|  | p | 0,218 | 0,547 | 0,207 | **0,039** | **0,005** | 0,723 |
| Random effects | N | 161 | 161 | 161 | 161 | 161 | 161 |
|  | R<sup>2</sup> Marginal/Conditional | 0,009 / 0,025 | - | 0,012 / 0,026 | - | - | - |

When analysing meiofaunal groups separately, group-specific patterns emerged: depth negatively influenced the taxonomic and phylogenetic richness of Platyhelminthes (GLMM: *χ*^2^=5.75, p=0.016; *χ*^2^=8.54, p=0.003, respectively) and negatively impacted the phylogenetic richness of Ostracoda (*χ*^2^=5.42, p=0.020) (Table 2 & Table 3). Mesh size significantly affected the taxonomic and phylogenetic richness of Gastrotricha (*χ*^2^=11.45, p=0.010; *χ*^2^=9.17, p=0.027), with the 50 µm retaining most of the richness (Figure 3; Table S2, Table S4). Habitat type significantly influenced the taxonomic diversity of Copepoda (*χ*^2^=12.08, p=0.034), Nematoda (*χ*^2^=12.85, p=0.025), Platyhelminthes (*χ*^2^=15.14, p=0.010) and Ostracoda (*χ*^2^=13.81, p=0.017) (Table 2; Table S3), as well as phylogenetic diversity of Platyhelminthes (*χ*^2^=11.69, p=0.039) and Ostracoda (*χ*^2^=16.68, p=0.005) (Table 3; Table S5) (Figure 4).

**Figure 3.**
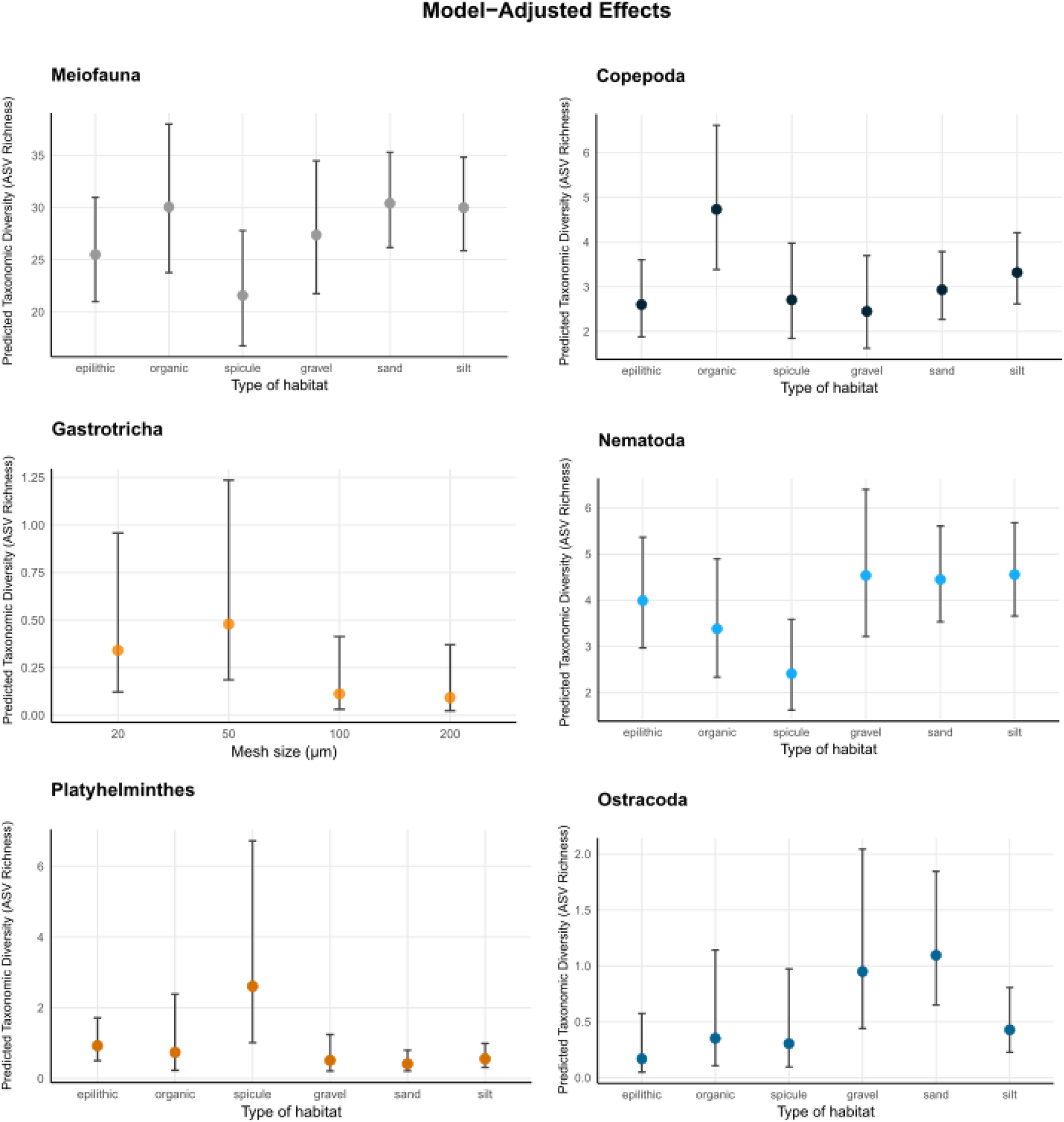
Predicted taxonomic alpha diversity (ASV richness) of the taxonomic groups that significantly vary across mesh sizes and habitat types. Lines and points represent the Estimated Marginal Means (predicted values) derived from the Generalized Linear Mixed Models (GLMMs), adjusted for sequencing depth and averaging across other covariates (habitat type and sampling depth). Vertical bars indicate the 95% Wald confidence intervals.

**Figure 4.**
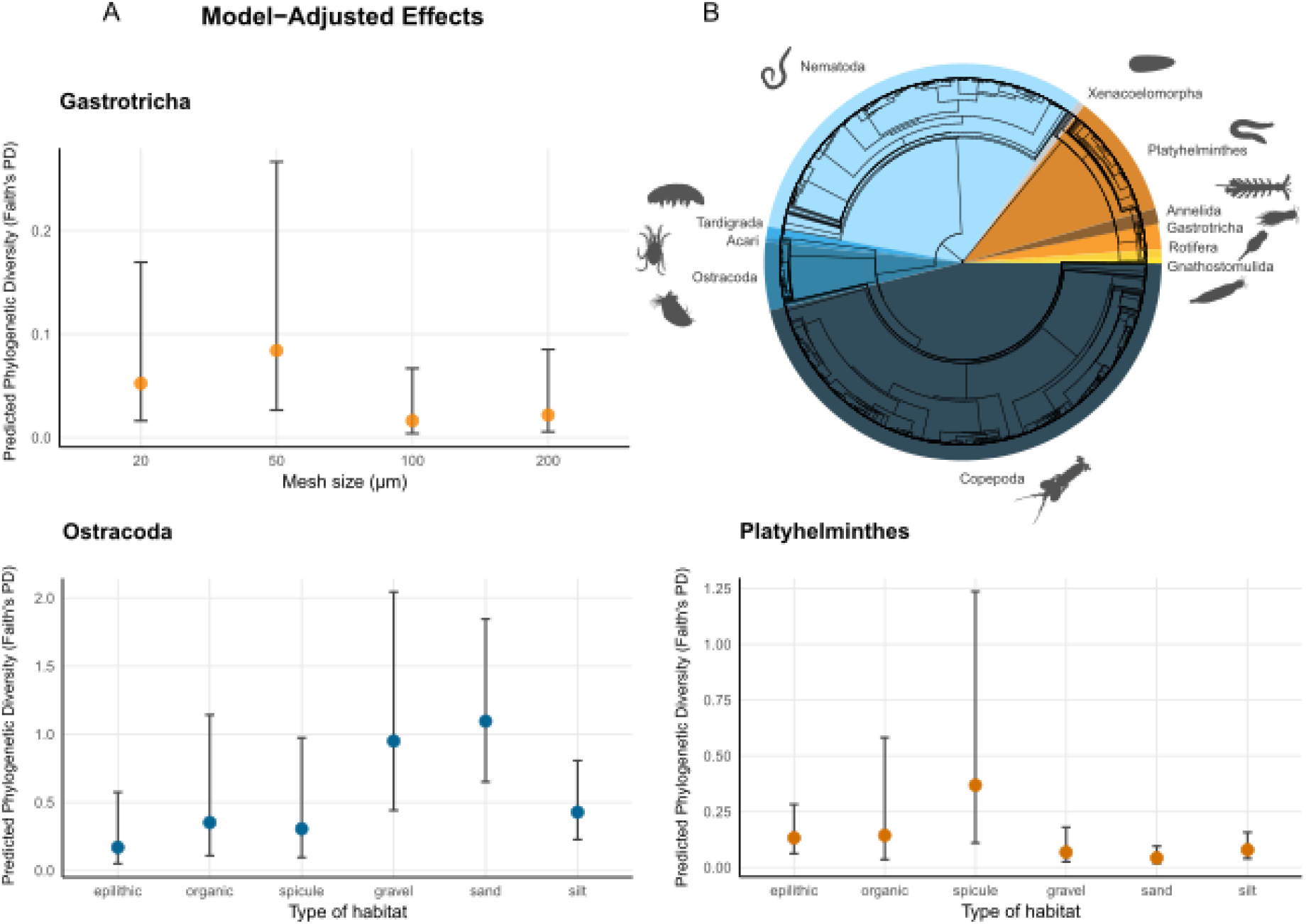
(A) Predicted phylogenetic richness across mesh sizes and habitat types for the total meiofaunal community and single taxonomic groups. Lines and points represent the Estimated Marginal Means (predicted values) derived from the Generalized Linear Mixed Models (GLMMs), adjusted for sequencing depth and averaging across other covariates (habitat type and depth). Vertical bars indicate the 95% Wald confidence intervals. Only significant results in the analyses are represented. (B) Neighbour Joining phylogenetic reconstruction of meiofaunal ASVs in our samples.

### Beta diversity: community composition

#### Taxonomic Beta Diversity

The overall meiofauna community composition was significantly influenced by all studied variables (Table 4). After controlling for sequencing effort, which explained a minor but significant proportion of the variance (PERMANOVA: R^2^=0.031, p<0.001), habitat type emerged as the primary driver of community structure (R^2^=0.13, p=0.001). This was followed by mesh size (R^2^=0.04, p=0.001) and depth, which accounted only for 1% of the variance (R^2^=0.01, p=0.001; Figure 5). At the group level, sequencing depth was significant for all taxa except Acari, with its explanatory power ranging from R^2^=0.009 in Annelida to R^2^=0.07 in Ostracoda. Habitat remained the most influential factor across almost all taxa, with its explanatory power ranging from 4% in Xenacoelomorpha to 39% in Acari (Table 4), although this effect was only statistically significant for Nematoda, Ostractoda, and Xenacoelomorpha. Mesh size explained a small but significant proportion of variance (all < 5%), except for the Ostracoda, Annelida, and Acari. Lastly, depth had a significant effect across all taxa except for Annelida (R^2^=0.01, p=0.453) and Acari (R^2^<0.01, p=0.519).

**Table 4.**
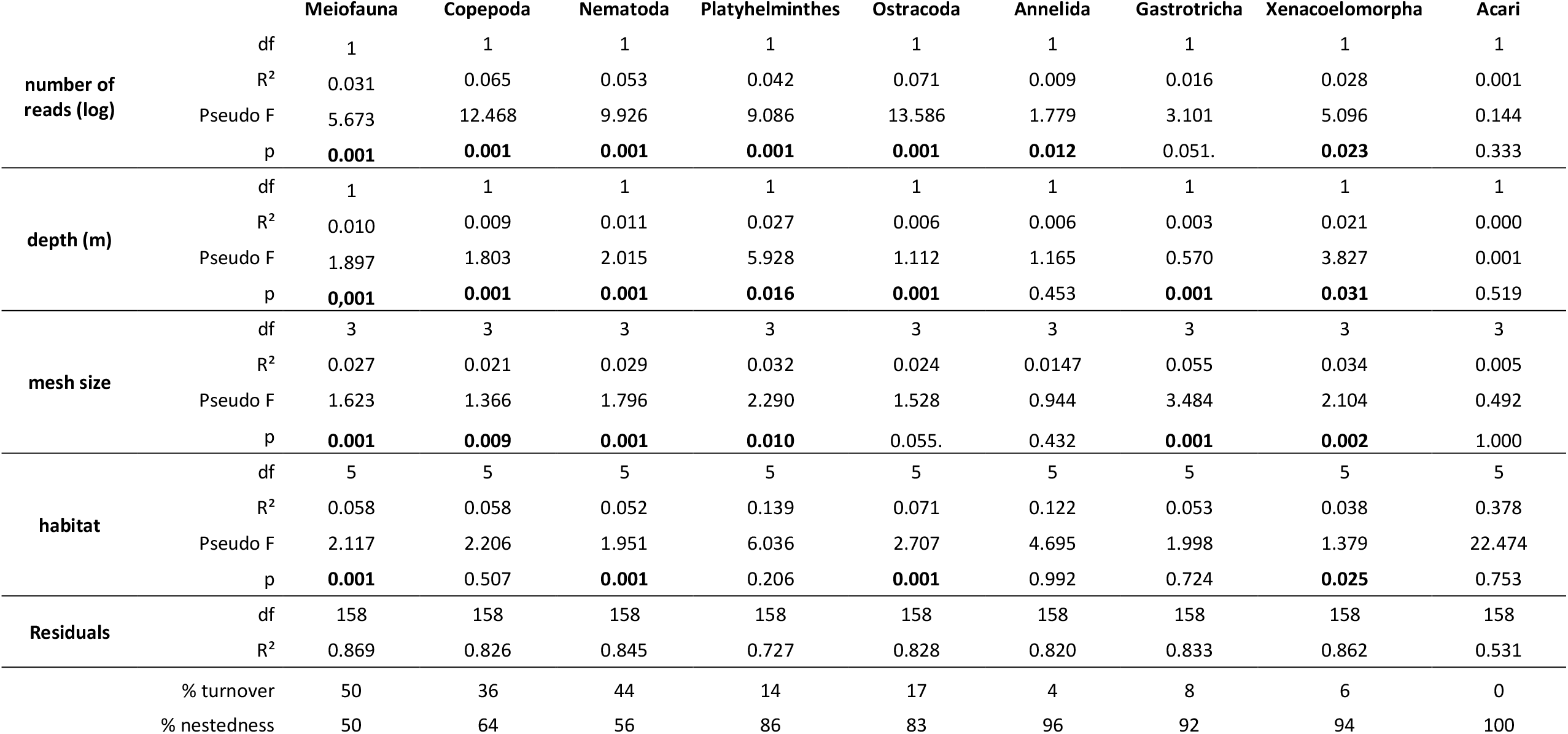
PERMANOVA results based on taxonomic beta diversity according to the explanatory variables: sampling depth, mesh size and type of habitat. The table shows the partitioning of variance for total taxonomic beta diversity (Jaccard index) and its components, turnover and nestedness.

**Figure 5.**
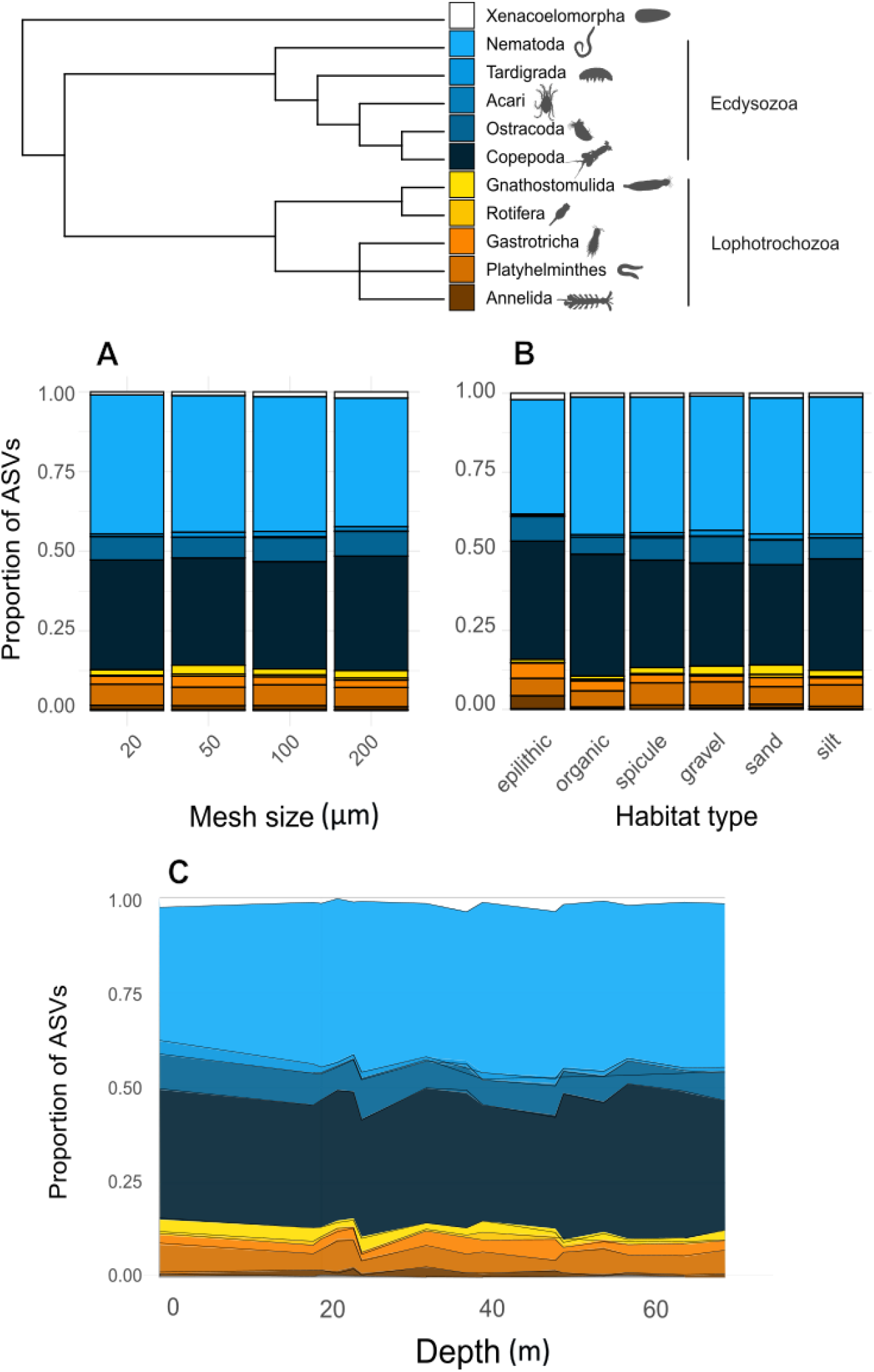
Taxonomic composition of the Antarctic meiofaunal community. Stacked bar plots (A, B) and area plots (C) represent the proportion of ASVs for each taxonomic group across sizes, habitat types and depth gradients. The cladogram at the top illustrates the phylogenetic relationships among studied groups, with topology based on Dunn et al. (2008).

Partitioning total beta diversity revealed that overall meiofaunal community shifts were equally distributed by turnover and nestedness (50% for both). However, at the level of individual taxonomic groups, community differences were primarily driven by nestedness, ranging from 64% in Copepoda to 100% in Acari (Table 4).

#### Phylogenetic Beta Diversity

Phylogenetic community structure was significantly influenced by all variables (Table 5). Sequencing effort accounted for a higher proportion of phylogenetic variance compared to taxonomic patterns (PERMANOVA: R^2^=0.10, p=0.001), explaining even more variance than habitat type (R^2^=0.05, p=0.002), although still relatively low. Lastly, mesh size (R^2^=0.03, p=0.001) and depth (R^2^= 0.01, p=0.001), explained a very low but significant variance.

**Table 5.** PERMANOVA results based on phylogenetic beta diversity according to the explanatory variables: sampling depth, mesh size and type of habitat. The table shows the partitioning of variance for total taxonomic beta diversity (based on PD-dissimilarity) and its components, turnover and nestedness.

|  |  | Meiofauna | Copepoda | Nematoda | Platyhelminthes | Ostracoda | Annelida | Gastrotricha | Xenacoelomorpha | Acari |
| --- | --- | --- | --- | --- | --- | --- | --- | --- | --- | --- |
| number of reads (log) | df | 1 | 1 | 1 | 1 | 1 | 1 | 1 | 1 | 1 |
|  | R<sup>2</sup> | 0.096 | 0.124 | 0.091 | 0.054 | 0.095 | 0.009 | 0.021 | 0.030 | 0.007 |
|  | Pseudo F | 19.233 | 26.922 | 17.886 | 11.728 | 18.699 | 1.862 | 4.000 | 5.586 | 1.684 |
|  | p | **0.001** | **0.001** | **0.001** | **0.001** | **0.001** | **0.006** | **0.042** | **0.018** | 0.114 |
| depth (m) | df | 1 | 1 | 1 | 1 | 1 | 1 | 1 | 1 | 1 |
|  | R<sup>2</sup> | 0.009 | 0.009 | 0.009 | 0.032 | 0.007 | 0.006 | 0.002 | 0.020 | 0.001 |
|  | Pseudo F | 1.839 | 1.944 | 1.713 | 7.090 | 1.482 | 1.182 | 0.349 | 3.696 | 0.281 |
|  | p | **0.001** | **0.001** | **0.001** | **0.019** | **0.001** | 0.401 | **0.003** | **0.009** | 0.310 |
| mesh size | df | 3 | 3 | 3 | 3 | 3 | 3 | 3 | 3 | 3 |
|  | R<sup>2</sup> | 0.027 | 0.021 | 0.030 | 0.029 | 0.021 | 0.013 | 0.061 | 0.035 | 0.013 |
|  | Pseudo F | 1.792 | 1.489 | 1.949 | 2.147 | 1.399 | 0.861 | 3.855 | 2.135 | 0.969 |
|  | p | **0.001** | 0.067 | **0.001** | **0.025** | 0.129 | 0.489 | **0.003** | **0.002** | 0.225 |
| habitat | df | 5 | 5 | 5 | 5 | 5 | 5 | 5 | 5 | 5 |
|  | R<sup>2</sup> | 0.048 | 0.046 | 0.045 | 0.122 | 0.075 | 0.123 | 0.036 | 0.037 | 0.270 |
|  | Pseudo F | 1.933 | 1.922 | 1.765 | 5.352 | 2.970 | 4.782 | 1.377 | 1.355 | 12.428 |
|  | p | **0.002** | 0.989 | **0.001** | 0.715 | **0.001** | 0.996 | 0.838 | **0.022** | **0.001** |
| Residuals | df | 158 | 158 | 158 | 158 | 158 | 158 | 158 | 158 | 158 |
|  | R<sup>2</sup> | 0.792 | 0.750 | 0.800 | 0.723 | 0.803 | 0.814 | 0.829 | 0.860 | 0.687 |
| % turnover |  | 30 | 16 | 29 | 7 | 6 | 1 | 4 | 4 | 0 |
| % nestedness |  | 70 | 84 | 71 | 93 | 94 | 99 | 96 | 96 | 100 |

At the group level, the influence of sequencing effort was most pronounced in Copepoda (R^2^ = 0.12) and Nematoda (R^2^= 0.09), while it remained non-significant for Acari. Habitat significantly affected the community composition of Nematoda (R^2^=0.04, p=0.001), Ostracoda (R^2^=0.07, p=0.001), Xenacoelomorpha (R^2^=0.04, p=0.022), and Acari (R^2^=0.27, p=0.001). Mesh size significantly affected Nematoda (R^2^=0.03, p=0.001), Platyhelminthes (R^2^=0.03, p=0.025), Gastrotricha (R^2^=0.06, p=0.003), and Xenacoelomorpha (R^2^=0.03, p=0.002). Depth affected the phylogenetic beta diversity of all taxa except for Annelida and Acari (Table 5).

The partitioning of phylogenetic beta diversity revealed that for the total meiofauna community, as well as for individual groups, nestedness was the primary driver of diversity shifts, ranging from 70 to 100% (Table 5).

### Degree of taxonomic novelty

The proportion of ASVs with less than 95% identity in GenBank in previous studies from different biogeographical areas ranged from 14% in the North Sea (Degenhart et al., 2021) to 73%, also in the North Sea (Mauffrey et al., 2020), with the previous study in Antarctica by Fonseca et al. (2017) having 30% (Figure S1, Table S6). On average, the proportions from the previous studies was 41.6±19.3; the probability that the proportion of 52% for our Antarctic survey would fall outside of the distribution is 0.590.

## DISCUSSION

### High meiofaunal diversity in Antarctica

Our results reveal a high diversity of marine meiofauna within the sediments of the Antarctic Ross Sea, identifying several potential species from 10 phyla with permanent meiofaunal representatives. These findings challenge the traditional perception of Antarctica and polar regions as poor and depauperate ecosystems compared to temperate and tropical latitudes (Rogers, 2007), a pattern that has been clearly confirmed for all freshwater and terrestrial biota in Antarctica, not only meiofauna (Pertierra et al., 2025). Diversity of Antarctic marine meiofauna is not poor: the structural complexity observed in our survey is not only comparable to, but in several cases exceeds, that reported for theoretically more productive or diverse biogeographical regions, for example, the diversity identified in our survey of Antarctic meiofauna surpasses what was found off the coast of Massachusetts (Polinski et al., 2019) and the Mediterranean Sea (Martínez et al., 2020) (Figure 2), although those comparisons are merely qualitative.

A comparison with the only previous study on marine meiofauna in Antarctica is needed: while Fonseca et al. (2017) reported a higher dominance of nematodes in Antarctica using 45 µm mesh, our multi-mesh approach demonstrated that the Antarctic benthos harbors a richer community when the full size-spectrum is accounted for (Table S1). Furthermore, Fonseca et al. (2017) used a different sequencing platform and sequencing depth, and focused exclusively on very shallow coastal sites (8-18 m) and found that they shared on average only 15% of OTUs with deep sea samples (290-515 m), suggesting a strong bathymetric zonation in the region. Since our sampling reached depths of up to 70 m, our study covers an intermediate shelf zone that might act as a transition between the shallow and deep-sea benthos.

Our beta diversity analyses demonstrated that mesh size may significantly influence the description of taxonomic composition (Table 4), suggesting that the choice of sampling mesh is not trivial. While a bigger mesh size might provide a coarse overview of the community, the 20 µm mesh is necessary for an exhaustive inventory of the meiofaunal community (Figure 6). This is particularly critical for taxa such as Copepoda, Nematoda, Platyhelminthes, Gastrotricha, and Xenacoelomorpha, whose community composition was significantly affected by mesh size (Table 4 & Table 5).

**Figure 6.**
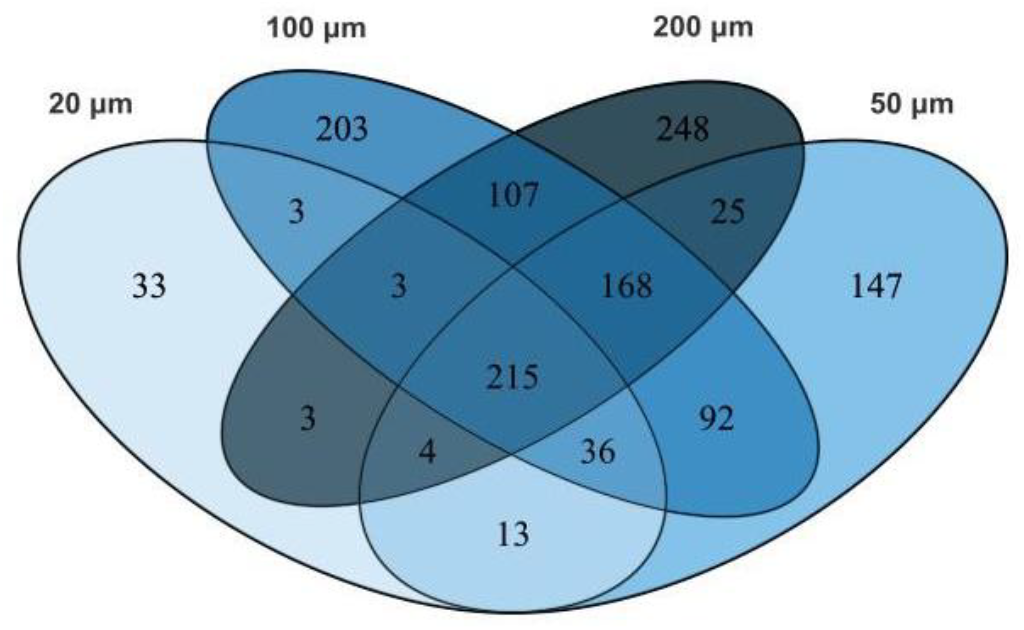
Venn Diagram showing number of unique ASVs for each mesh size and the number of shared ASVs across different mesh sizes.

### Biogeography and taxonomic resolution

A striking finding of our analysis is the detection of ASVs that are 100% identical to those found in geographically distant regions, such as the North Sea and the North Atlantic (Atherton & Jondelius, 2020; Haenel et al., 2017; Polinski et al., 2019). Historically, the apparent cosmopolitan distribution of microscopic marine animals that lack dispersal stages has been used to illustrate the “meiofauna paradox” (Giere, 2009) and the “everything-is-everywhere hypothesis” (Fenchel & Finlay, 2004). However, in our survey we could suggest that the shared sequences are likely an artifact of the conservative nature of the 18S rRNA gene marker rather than evidence of global dispersal. The 18S rRNA gene is known for its evolutionary stasis in several meiofaunal groups (Tang et al., 2012); thus, geographic isolation may have led to speciation events that remain undetected at the level of 18S analyses.

Beyond the potential underestimation of species-level endemism, our dataset reveals that the level of potentially unexplored diversity at higher taxonomic levels (less than 95% identity in GenBank) seems to be comparable to that of any other metabarcoding survey of the meiofauna in other biogeographical regions. Thus, of course several endemic species are expected to be discovered in the meiofauna in Antarctica, but this is similar to what happens for any other biogeographical area (e.g. Curini-Galletti et al., 2012; Martinez et al., 2019). Yet, no exclusive radiation of higher groups can be expected, contrary to what happened in the Antarctic marine habitats at least for fish (Matschiner et al., 2011; Di Prisco et al., 2012), echinoderms (Moles et al., 2015), isopods (Lecointre et al., 2013), amphipods and sponges (Bowen et al., 2020).

### Habitat heterogeneity as the primary driver of community structure

Beyond the geographical scale, our study demonstrates that local habitat type, measured as the type of substrate, is the most influential factor determining community composition, expressed as taxonomic and phylogenetic beta diversity, as it is commonly expected for any meiofaunal communities in other parts of the world (Martínez et al., 2021; Matsko et al., 2026; Mirto et al., 2010). Since community shifts were mainly driven by nestedness in most groups, our findings suggest that instead of a complete replacement of species between habitats, certain substrates support a richer subset of the community while others act as environmental filters (Gansfort et al., 2020).

This habitat-driven filtering was also reflected in alpha diversity patterns. Among taxa significantly influenced by the type of habitat, the ones harboring the highest richness were group-specific: organic sediments for Copepoda, gravel and sand for Ostracoda and spicule mats for Platyhelminthes (Figure 3). This could suggest an adaptation of the groups specific to those habitats (Ingels & Zeppilli, 2023).

## CONCLUSIONS

Our biodiversity survey through metabarcoding revealed that, contrary to terrestrial and freshwater habitats in Antarctica (Pertierra et al., 2025), diversity in marine Antarctic meiofauna, despite the apparent simplicity of Antarctic marine environments (Rogers, 2007) and regardless of the potentially harsh environmental conditions of the Antarctic coastal habitats, is not poor and depauperate but is as rich and diverse as in other biogeographical regions.

